# Generative design of intrinsically disordered protein regions with IDiom

**DOI:** 10.64898/2026.04.10.717777

**Authors:** Jason X. Liu, Sebastian Ibarraran, Frank Hu, Abigail Park, Alexander R. Dunn, Grant M. Rotskoff

**Affiliations:** Department of Chemical Engineering, Stanford University, Stanford, CA 94305; Department of Chemistry, Stanford University, Stanford, CA 94305; Institute for Computational and Mathematical Engineering, Stanford University, Stanford, CA 94305

## Abstract

Intrinsically disordered protein regions are ubiquitous across all kingdoms of life. These structurally heterogeneous regions play central roles in cellular processes such as transcriptional regulation, cellular signaling, and subcellular organization, yet they have remained largely inaccessible to rational design. Structure-based generative methods are not applicable to proteins that lack a stable fold, and existing sequence-based approaches for disordered regions rely on sampling methods that do not capture the evolutionary statistics of natural disordered regions. Here, we introduce IDiom, an autoregressive protein language model trained on 37 million intrinsically disordered region sequences curated from the AlphaFold Database. Trained using a fill-in-the-middle data augmentation, IDiom generates disordered region sequences conditioned on their surrounding structured context, as well as fully disordered proteins without any context. The model generates diverse sequences that recapitulate biologically relevant sequence features of natural disordered regions, and we demonstrate that post-training via reinforcement learning with a subcellular localization reward model produces sequences with features which are consistent with known sequence determinants of compartment-specific localization. These results establish IDiom as a general platform for the generative design of intrinsically disordered proteins and regions.

## Introduction

Intrinsically disordered protein regions are ubiquitous across all kingdoms of life. The functional relevance of intrinsically disordered regions (IDRs) has become increasingly clear recently, in spite of the classical dogma of protein biology that structure implies function [1]. While IDRs do not adopt well-defined folds, these sequences can act as flexible linkers [2], multivalent signaling hubs [3], and drivers of biomolecular condensate formation [4, 5, 6], playing crucial roles in biological processes such as transcriptional regulation, chromatin organization, and subcellular organization [1, 7].

Rational design of IDRs would unlock new functional controls in bioengineering, including tunable condensate phase behavior, precise regulation of cell signaling, and targeted protein localization [8]. While much of the recent progress in protein design has been driven by accurate structure prediction [9, 10] and diffusion-based generative models [11, 12, 13], direct application of these approaches to IDRs is untenable due to their lack of a stable folded structure.

Sequence-based generative models have also been extensively explored recently for applications in protein design. Transformer-based [14] protein language models (PLMs) have been trained on corpora of full length protein sequences from databases such as the UniProt Reference Clusters and the Big Fantastic Database. These models learn rich evolutionary statistics over amino acid sequences and have enabled the design of novel proteins and functional variants [15, 16, 17, 18, 19, 20]. Nevertheless, the design of intrinsically disordered regions with existing PLMs is not straight-forward: since structured domains outnumber intrinsically disordered regions in the sequence databases on which current PLMs are trained, the generative prior is largely biased towards folded domains [15]. Alternative sequence-based approaches to IDR design have attempted to use sampling-based methods to construct IDRs from compositional rules or simple statistical models [21, 22, 23, 24]. However, these approaches cannot be conditioned on any surrounding sequence context of generated IDRs, and they do not capture the evolutionary statistics which emerge from training on large corpora of natural protein sequences.

Here, we address this gap by training a 122M parameter autoregressive, decoder-only protein language model called IDiom using a dataset of 37 million intrinsically disordered regions curated from the AlphaFold Database (Figure 1a) [25, 26]. We apply a fill-in-the-middle transformation to the training data [27, 28] to enable the model to generate IDR spans conditioned on their surrounding context, a capability essential for the design of disordered regions [8]. Training on this dataset of disordered sequences allows IDiom to learn a generative prior over the sequence statistics of natural disordered regions, and we demonstrate that the model generates diverse sequences that recapitulate the compositions, patterning, and motifs of natural intrinsically disordered regions. We further show that IDiom learns in-context: given the flanking sequence context of a specific protein, the model generates disordered spans whose sequence features are more appropriate for that context than unprompted generations. Finally, we demonstrate that IDiom can be post-trained using reinforcement learning, and we apply this to design disordered sequences with targeted subcellular localization [23]. Together, these results establish IDiom as a general platform for the generative design of intrinsically disordered proteins and regions.

**Fig. 1:**
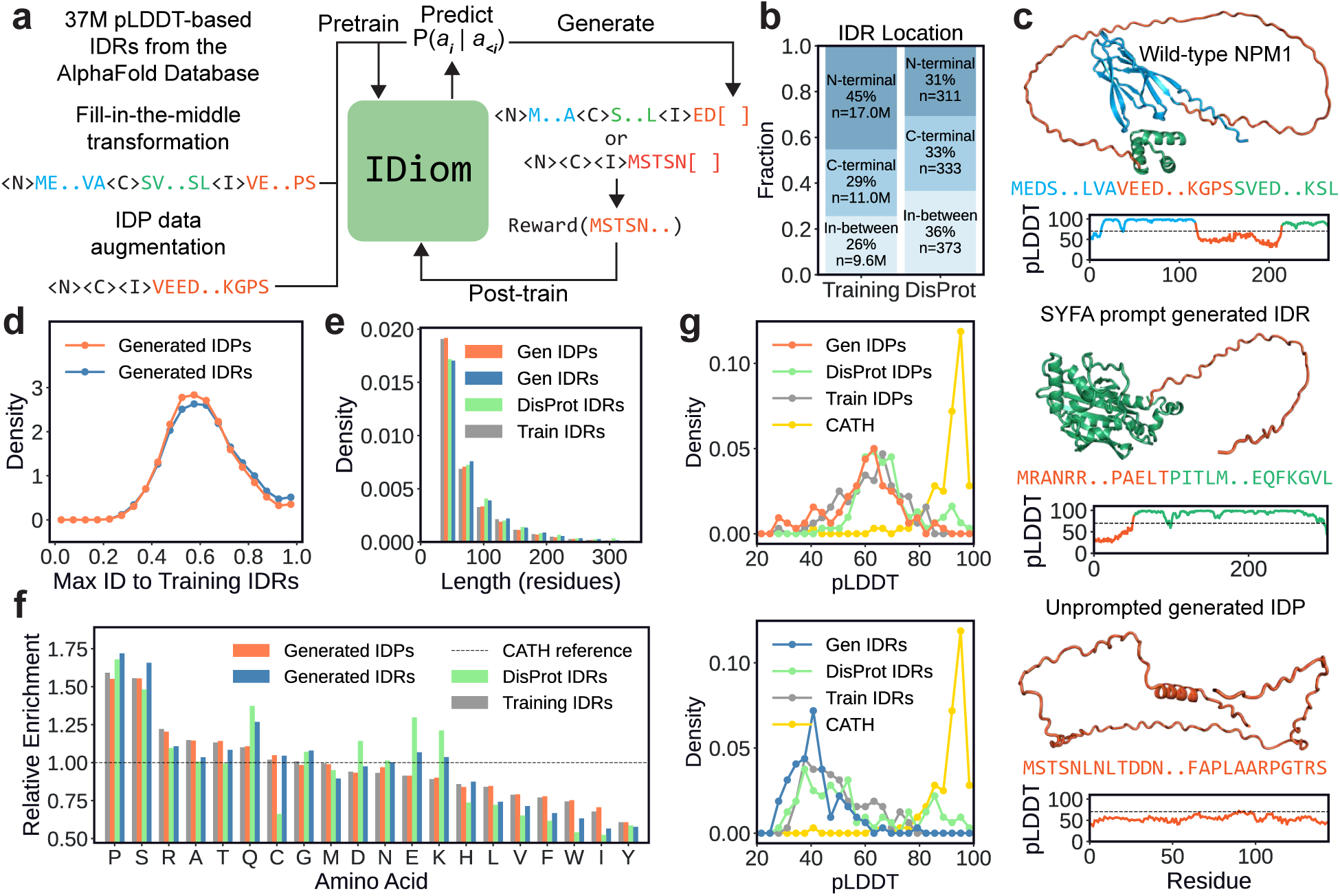
Data curation, training, and generative modeling of intrinsically disordered regions (IDRs) and proteins (IDPs) using IDiom. **(a)** Schematic depicting data preparation, pre-training, sequence generation, and post-training of IDiom. **(b)** Distribution of the sequence locations of 37M IDRs curated from the AlphaFold Database (AFDB), as well as the locations of experimentally validated DisProt IDRs. **(c)** AlphaFold2 (AF2)-predicted structures, amino acid sequences, and plots of predicted local distance difference test (pLDDT) values of an example IDR curated from AFDB (NPM1, upper), an IDR generated by IDiom using SYFA as the prompt (middle), and an unprompted generated IDP (lower). Blue and green regions correspond to the N-terminal and C-terminal flanking context around the IDR, which is orange. **(d)** Distribution of maximum sequence identities of unprompted generated IDPs and DisProt-prompt generated IDRs relative to the 37M IDRs of the training set, calculated using MMseqs2. **(e)** Distribution of sequence lengths of unprompted generated IDPs, DisProt-prompt generated IDRs, natural DisProt IDRs, and training set IDRs. **(f)** Relative enrichment of amino acid compositions of unprompted generated IDPs, DisProt-prompt generated IDRs, natural DisProt IDRs, and training set IDRs. The horizontal dashed line is the reference composition of folded CATH domain sequences. **(g)** (Upper): AF2 prediction pLDDTs of generated unprompted IDPs, natural DisProt IDRs removed from their surrounding context (DisProt IDPs), training set IDRs removed from their surrounding context (train IDPs), and CATH domains. (Lower): AF2 prediction pLDDTs of DisProt-prompt generated IDRs, natural DisProt IDRs within their context, training set IDRs within their context, and CATH domains.

## Results

### Protein language modeling for intrinsically disordered proteins and regions

To curate a dataset of intrinsically disordered region (IDR) sequences for model training, we first use AlphaFold2 (AF2) predicted structures from the AlphaFold Database (AFDB) [26] to identify IDRs of proteins. We use low AF2 predicted local distance difference test (pLDDT) values as a predictor of disorder, as this has been demonstrated to correlate strongly with experimental measurements of disorder [29, 30, 31]. To curate these IDRs from the database, we first cluster AFDB sequences at 90% sequence identity before applying a windowed pLDDT-based threshold to identify IDRs [32], with proteins containing multiple IDRs contributing multiple records to the dataset. Finally, we discard IDRs shorter than 30 residues, IDRs which reside in proteins whose full length is greater than 512 residues, and proteins whose entire length is low-pLDDT. This process yields a dataset of 37M IDRs and their positions within their associated full length proteins (see Methods for more details). Figure 1c (upper) depicts an example low-pLDDT IDR, in orange, extracted by this process for the human protein NPM1 (UniProt: P06748).

We use two baselines to validate our data curation strategy. First, we use 1,017 experimentally validated IDRs from the DisProt database as a ground-truth dataset of IDRs [33]. Second, we use 1,000 randomly chosen sequences from the Class, Architecture, Topology, and Homologous superfamily (CATH) database clustered at 60% identity (S60) as a ground-truth dataset of protein domains with well-defined folded structures [34]. Using secondary structure content calculations, we show that this method extracts IDRs with substantially lower secondary structural content compared to CATH sequences (Figure S1). We note that our analysis shows that AF2 assigns high secondary structural content (with low confidence) to a fraction of the extracted IDRs, although this effect is also reflected in the set of DisProt IDRs. We additionally run orthogonal disorder predictions on a subset of the dataset and further validate that the curated sequences score similarly to DisProt sequences on these metrics (Figures S2, S3).

Intrinsically disordered regions are naturally located at various locations within a protein sequence, with approximately 45% of the AFDB-curated IDRs located at the N-terminus, 29% at the C-terminus, and 26% in between other regions of the protein; similar percentages are observed for experimentally validated IDRs from the DisProt set (Figure 1b). To enable IDiom to infill sequences of intrinsically disordered regions at arbitrary locations within a protein, we employ a fill-in-the-middle data transformation [27, 28]. In this approach, we prepend the special token <N> to residues in the N-terminal flanking context before the IDR, the token <C> to residues in the C-terminal context after the IDR, and the token to residues of the IDR span itself. Then, we transform the sequence by relocating and the IDR span to the end of the entire sequence, thus allowing the standard causal language modeling objective to generate IDR spans conditioned on any preceding N-terminal and succeeding C-terminal flanking context (see Figure 1a). In addition, to enable the generation of intrinsically disordered proteins (IDPs) with no surrounding context, we augment the dataset by creating records in which take each curated IDR and delete their N- and C-terminal flanking context, enabling unconditioned generation. The final dataset comprises 74M sequences in total (37M IDRs and 37M IDPs), which we use to pre-train IDiom. Additional details of the data augmentation, model architecture, and training procedure are provided in the Methods section.

### IDiom generates diverse disordered regions and proteins

To characterize the pre-trained model, we first generate two sets of sequences: 100,000 unprompted IDPs (referred to as generated IDPs), and a set of context-prompted IDRs in which 100 IDRs are generated for each of 1,017 experimentally validated IDRs from the DisProt set, using the IDR flanking contexts as prompts (referred to as generated IDRs). Representative examples of a DisProt context-prompted IDR and an unprompted IDP are shown in Figure 1c (middle) and 1c (lower), respectively.

We find that across multiple metrics, IDiom generates sequences that closely resemble natural DisProt IDRs while remaining diverse and distinct from the training data. The distribution of maximum sequence identities to training set IDRs peaks broadly around 60%, indicating that most generated sequences are substantially dissimilar to any sequence seen during training (Figure 1d). Generated IDR and IDP lengths are also consistent with those of both the training set and DisProt IDRs, with most sequences below 100 residues long and a tail extending to approximately 300 residues long (Figure 1e).

To analyze the compositional biases of the IDRs and IDPs, we compute the amino acid enrichment of training, generated, and DisProt sequences relative to a baseline composition of the folded CATH domains (Figure 1f). Consistent with established compositional biases of disordered regions [35, 36], generated sequences are strongly enriched in proline and serine, and depleted in order-promoting aliphatics such as leucine, isoleucine, and valine, as well as the aromatics phenylalanine, tryptophan, and tyrosine. The trends for training and generated sequences closely match natural DisProt IDRs across most amino acids, with the closest agreement observed for IDRs generated with the DisProt flanking contexts as prompts. We note that the discrepancies with DisProt in the generated compositions of cysteine and charged residues (glutamic acid, lysine, and aspartic acid), may be attributed to the small size and selection bias of the DisProt dataset.

We next assess the extent to which the generated sequences are disordered by predicting their structures using ColabFold [37]; the resulting pLDDT distributions are shown in Figure 1g. We run the structure predictions in two settings. In the first, we compare the unprompted generated IDPs to DisProt and training IDRs which are provided to ColabFold as standalone sequences, without their flanking context (i.e. as IDPs). In this setting, the pLDDT for a given sequence is averaged across the entire IDP, generated or extracted (Figure 1g (upper)). In the second, we compare the DisProt-prompt generated IDRs to the full protein DisProt and training sequences. In this setting, we provide the full generated, training, or DisProt sequence to ColabFold, and the pLDDT is only averaged across the IDR span (Figure 1g (lower)). Across both settings, generated sequences exhibit pLDDT distributions that closely mirror those of DisProt as well as the training set, confirming that IDiom is able to generate sequences which are predicted to be disordered to the same extent as natural IDRs. We note that the pLDDTs of IDPs in isolation are higher than for IDRs within their flanking context, which may be due to dataset biases in the AF2 training data. We additionally calculate secondary structure metrics and conduct orthogonal disorder predictions on these generated sequences, and we find that the metrics compare similarly to DisProt IDRs (Figures S1–S3).

### Generated sequences capture the residue patterning of natural disordered regions

IDRs exhibit sequence patterning features that differ substantially from those of folded domains, including characteristic charge distributions, hydrophobic residue patterning, and low-complexity compositions [38, 36, 39]. Figure 2a shows representative generated sequences which illustrate canonical IDR features such as Q/N-rich low-complexity regions [40], polyampholyte charge block patterning [41], prion-like aromatic/glycine patterning [4], and proline enrichment [35]. To quantify how well IDiom recapitulates these properties, we computed the distributions of several sequence-level metrics for the generated, training, and DisProt disordered sequences, as well as the folded CATH domain sequences (Figure 2b-e).

**Fig. 2:**
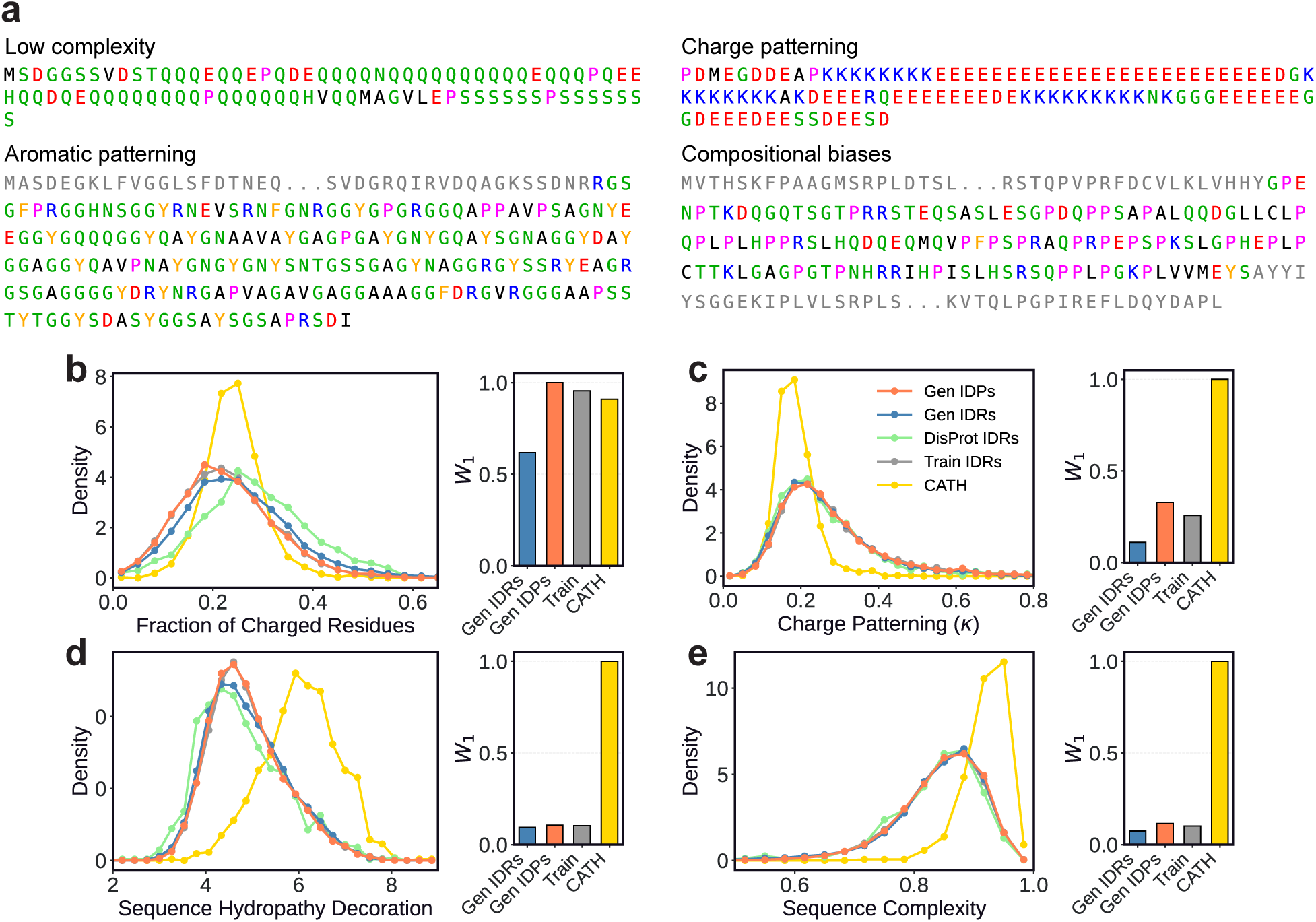
IDiom generates intrinsically disordered regions and proteins which capture the sequence patterning features of natural sequences. **(a)** Example IDPs (upper row) and IDRs (lower row) generated using IDiom which exhibit canonical sequence features of intrinsically disordered regions, including low complexity regions, charge blockiness, aromatic patterning, and compositional biases. Greyed-out residues correspond to the flanking context of the IDR. **(b)–(e)** Distributions of various sequence metrics for generated IDPs and IDRs, training set IDRs, DisProt IDRs, and folded CATH domains. Right subplots: Normalized Wasserstein-1 (*W*_1_) distance between the distributions of DisProt IDRs and all other distributions. **(b)** Fraction of charged residues (FCR). **(c)** Charge patterning *κ* parameter. **(d)** Sequence hydropathy decoration (SHD). **(e)** Sequence complexity quantified by the SEG algorithm.

Electrostatic interactions strongly influence the conformational behavior of IDRs, and both the overall charge content and its linear patterning are closely linked to physical properties and biological function [42, 43, 41, 39]. The fraction of charged residues (FCR, Figure 2b) distinguishes strongly charged polyampholytes from weakly charged sequences. We find that the generated and natural IDRs and IDPs span a wider range of FCR values than folded CATH domains, reflecting the heterogeneity of natural IDRs, which range from highly charged sequences in which electrostatic repulsion drives disorder, to weakly charged low-complexity sequences [44].

To characterize charge patterning, we calculate the linear charge patterning parameter *κ*, which quantifies the deviation of a given sequence from a maximally charge segregated permutation of the same sequence [44]. *κ* ≈ 0 indicates well-mixed opposite charges and *κ* ≈ 1 indicates segregation into blocks of the same charge, and we plot the distributions of these values in Figure 2c. Consistent with prior work linking charge segregation in IDRs to intermolecular interactions and phase behavior [41, 45], natural IDRs from the training and DisProt sets exhibit a tail toward high *κ* values relative to CATH domains, a feature that IDiom reproduces with its generated sequences.

We next examined hydrophobic patterning using the sequence hydropathy decoration (SHD) metric, which quantifies the spatial clustering of hydrophobic residues along the chain [46]. Generated sequences exhibit substantially lower SHD values than folded CATH domains (Figure 2d), consistent with the reduced hydrophobic clustering in disordered regions that prevents hydrophobic collapse [46]. This trend closely matches natural DisProt and training set IDRs, and it contrasts with folded CATH domains, which exhibit higher SHD values, reflecting the locally concentrated hydrophobic residues required to stabilize buried protein cores [47].

Finally, we assessed sequence complexity using the SEG algorithm, which computes the average compositional entropy over a sliding window [48]. IDRs frequently contain low-complexity segments [49], and natural DisProt and training set IDRs show lower complexity than folded CATH domains. Generated IDRs and IDPs closely reproduce this shift, with the complexities of generated sequences matching the DisProt distribution well (Figure 2e). All together, these results demonstrate that IDiom has learned the sequence grammar of disordered regions across multiple metrics, and that generated sequences closely recapitulate the sequence patterning features of natural IDRs.

### Conditioned generation recapitulates context-specific IDR sequence features

The previous analysis demonstrates that IDiom can generate sequences with patterning features and biophysical properties which are similar to natural IDRs. To quantify the agreement between generated and natural IDRs, we compute the normalized Wasserstein-1 distance (*W*_1_) [50] between the distributions of each sequence metric and the DisProt reference distribution (right subplots of Figure 2b-e). Across most metrics, *W*_1_ distances for generated sequences are low relative to folded CATH domains, confirming that IDiom produces sequences that are statistically much closer to natural IDRs than to folded proteins. We find that in Figure 2b, the shift in mean values for generated and training sequences relative to DisProt leads to relatively larger *W*_1_ values compared to the other metrics we consider, but we note that the shift relative to the training data likely results from the relatively small set of IDRs in DisProt. We also note that the shape of the broad distribution of FCR values for generated and training sequences matches that of DisProt more closely than the narrower CATH distribution. Furthermore, *W*_1_ distances for DisProt context-prompted IDRs are consistently lower than for unprompted IDPs across all metrics, demonstrating that conditioning on flanking sequence context shifts IDiom’s generations toward the sequence features of the natural IDRs that reside inside those contexts.

To further illustrate the model’s ability to learn via this in-context conditioning, we consider the human protein NPM1 (UniProt: P06748) as a case study. NPM1 has an IDR (residues 119–242) which drives nucleolar phase separation through charge block patterning that mediates interactions with itself [41] as well as binding partners such as SURF6 [51]. Using the flanking regions around this IDR as the prompt (green and blue domains in Figure 1c (upper)), we generated 100,000 IDRs and filtered out any with sequence identity *>* 90% to the wild-type (WT) NPM1 IDR (Figure 3a). Figure 3b shows the distribution of *κ* values for NPM1-prompted generations, randomly scrambled versions of those sequences, and the dataset of DisProt IDRs. We find that the *κ* distribution of NPM1-prompted generations is peaked near the WT NPM1 value, and is substantially shifted toward higher charge segregation than either randomly scrambled sequences or generic DisProt IDRs, indicating that the model has learned to generate sequences with substantial charge block patterning, when conditioned on the NPM1 flanking contexts.

**Fig. 3:**
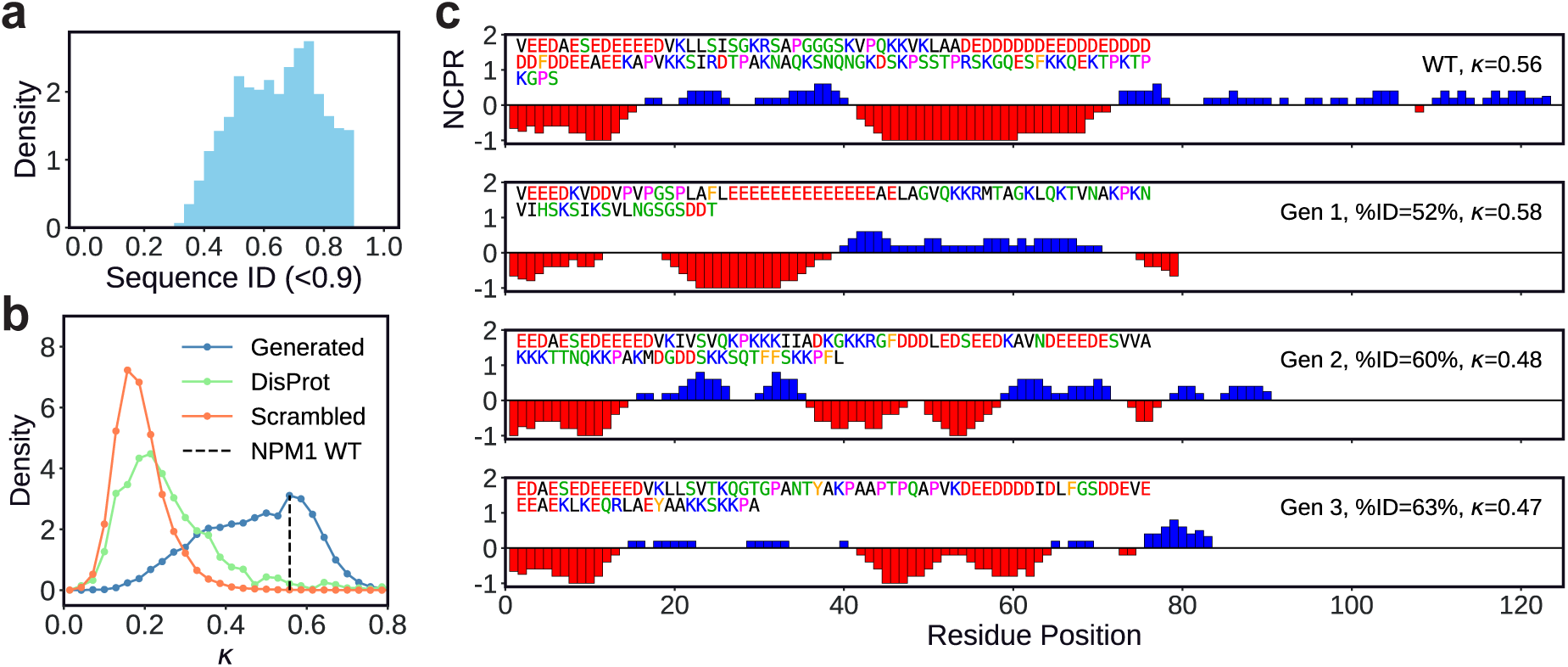
Conditioned generation and in-context learning enables generation of disordered regions which capture biologically relevant sequence features. **(a)** Distribution of sequence identities between the wild-type (WT) NPM1 IDR and sequences generated with the NPM1 IDR’s flanking context as the prompt. Generated sequences with identities greater than 0.9 are filtered out. **(b)** Distribution of *κ* values for the WT NPM1 IDR (black vertical dashed line), IDRs generated with the NPM1 context as prompt (blue), randomly scrambled versions of the generated IDRs (orange), and DisProt IDRs (green). **(c)** Plots of the net charge per residue along the sequence. The WT NPM1 IDR is in the upper row, and three representative generated IDRs are below. The sequence identities relative to the WT, *κ* values, and the amino acid sequences themselves are printed within the plots.

Figure 3c shows linear plots of net charge per residue (NCPR) for the WT NPM1 IDR and representative generated sequences with high *κ* values but low sequence identity to the WT IDR. These plots illustrate that IDiom generates sequences with alternating blocks of positive and negative charge which closely mirror the architecture of the WT IDR, despite sharing little sequence identity with it. Together, these results demonstrate that IDiom is able to use flanking context to reproduce the biologically relevant sequence features of a given IDR, and that the model is able to generate diverse sequences that preserve the key sequence features that underlie biological function.

### Sequence optimization using reinforcement learning

Our results demonstrate that IDiom has learned a strong generative prior over IDR sequence space, and that the model produces diverse and biologically realistic sequences. This naturally positions the pre-trained model as a starting point for sequence design: post-training IDiom with external rewards allows us to steer generation towards sequences that score well on specified objectives while remaining IDR-like. Unlike supervised fine-tuning on labeled datasets, post-training via reinforcement learning allows us to optimize arbitrary reward functions such as computational predictors and reward models trained on experimental data. Reinforcement learning can additionally incorporate explicit regularization to control sequence diversity, length, and deviation from the base model [52, 53, 54].

As a design target, we focus on subcellular localization, as the ability to engineer protein localization could enable both targeted delivery of therapeutics and modulation of synthetic condensates [55, 56]. As the reward model, we use ProtGPS, a neural network trained with ESM2 embeddings to predict the probability of a given protein sequence localizing to each of twelve specific subcellular compartments [23, 16]. Here, we post-train IDiom to optimize localization to four compartments: the nucleolus, chromosomes/chromatin, P-bodies, and stress granules. These compartments were chosen because they are known to be enriched in proteins with IDR-specific sequence features, including charge segregation, RNA-interacting motifs, nuclear localization signals, and post-translational modification sites, thus providing a clear test of whether RL post-training can induce the generation of biologically relevant sequence features.

We optimize IDiom using the reinforcement learning algorithm Group Relative Policy Optimization (GRPO) [57, 58], and we generate unprompted IDPs in all post-training runs (overview in Figure 4a). To prevent reward hacking and to maintain sequence diversity, we incorporate three forms of regularization: a Kullback-Liebler (KL) divergence penalty *D*_KL_ to prevent excess divergence of the post-trained model from the pre-trained base model, a quadratic penalty around a target Shannon entropy of *H* = 2.7 nats to prevent diversity collapse, and a quadratic penalty around a target sequence length of 100 residues. Figure 4b shows training curves for the ProtGPS reward, *D*_KL_, and mean sequence length versus post-training optimizer steps. The ProtGPS reward increases steadily across all four compartments, the mean sequence lengths converge to the target value, and *D*_KL_ remains below 0.4 throughout training, confirming that post-training successfully optimizes the reward without diverging substantially from the pre-trained base distribution.

**Fig. 4:**
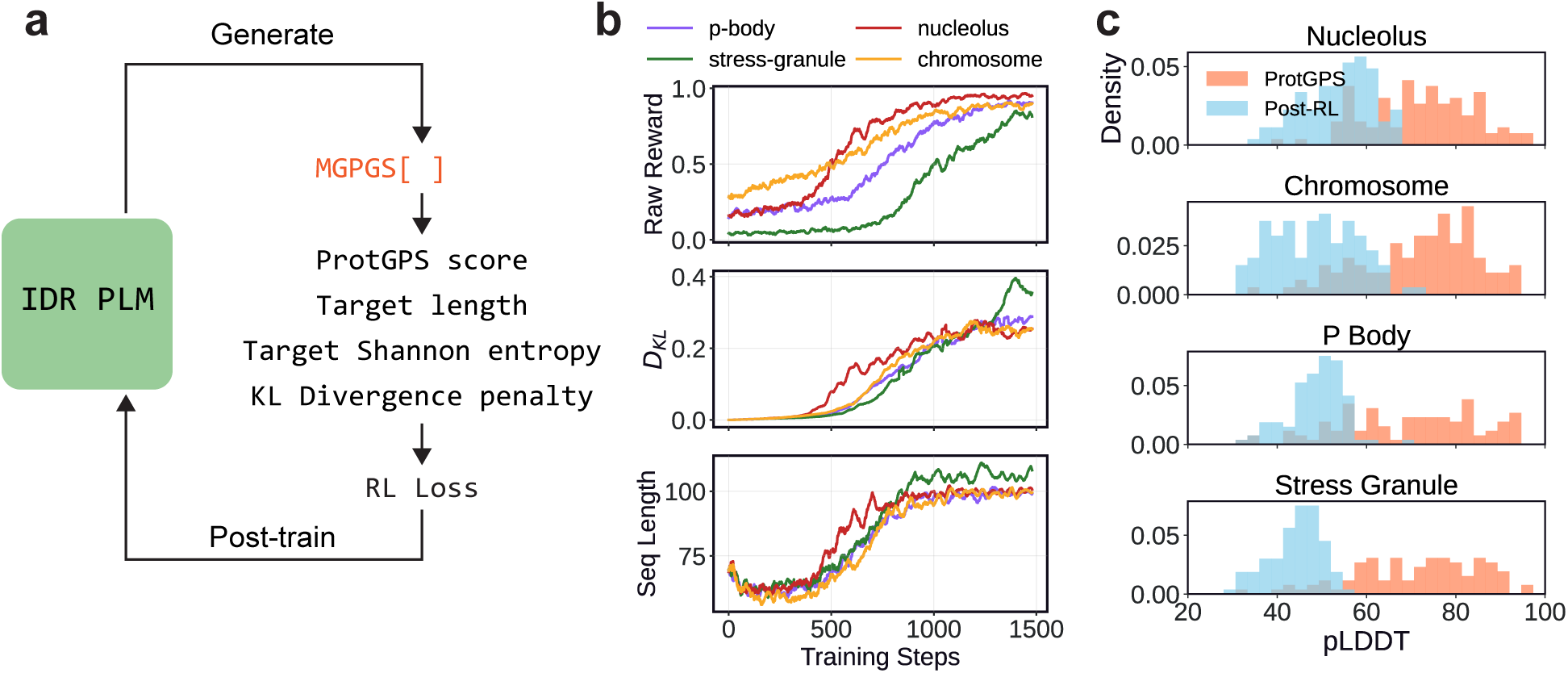
Post-training IDiom with reinforcement learning enables the generation of sequences aligned with an external reward model. **(a)** Schematic depicting the reinforcement learning process for post-training IDiom. ProtGPS is a reward model which returns a score between 0 and 1 indicating the probability of a given sequence localizing to a chosen subcellular compartment. Quadratic penalties are applied to sequences which deviate from the target ProtGPS score, target length, and target Shannon entropy, and a penalty is applied for an increased Kullback-Liebler (KL) divergence of the post-trained model from the pre-trained base model. These metrics are combined in a reinforcement learning loss which is used to update the IDiom policy. **(b)** Training curves depicting the ProtGPS score for a given compartment, magnitude of the KL-divergence, *D_KL_*, and the average generated sequence length, as a function of post-training optimizer steps. **(c)** Distribution of AlphaFold2 pLDDTs of sequences generated from IDiom checkpoints after 1500 post-training optimizer steps, as well as pLDDTs of the original sequences used to train the ProtGPS reward model.

Figure 4c shows the distributions of AlphaFold2 pLDDT values for sequences generated from each post-trained checkpoint, alongside the pLDDT distribution of the full length proteins used to train the ProtGPS predictor. The ProtGPS training sequences have a wide distribution of pLDDTs, with a substantial fraction of high-pLDDT residues, reflecting the fact that ProtGPS was trained on full length proteins containing both folded domains and disordered regions. However, sequences generated by the post-trained IDiom models maintain low pLDDT values, comparable to those of natural IDRs, demonstrating that the KL regularization successfully prevents the model from drifting towards the sequence features of folded proteins in the ProtGPS training set. This further confirms that post-training steers generation towards the desired compartment localization while preserving the disordered nature of generated sequences.

### Post-trained models generate sequences with compartment-specific features

After post-training with the ProtGPS reward, we generate 10,000 sequences from each localization-optimized checkpoint and analyze their amino acid compositions. Sequences targeting specific subcellular compartments are expected to have compositional biases that reflect their local biochemical environments, and we find that post-trained generations exhibit biologically interpretable compositional shifts relative to both base model generations and DisProt IDRs (Figure 5).

**Fig. 5:**
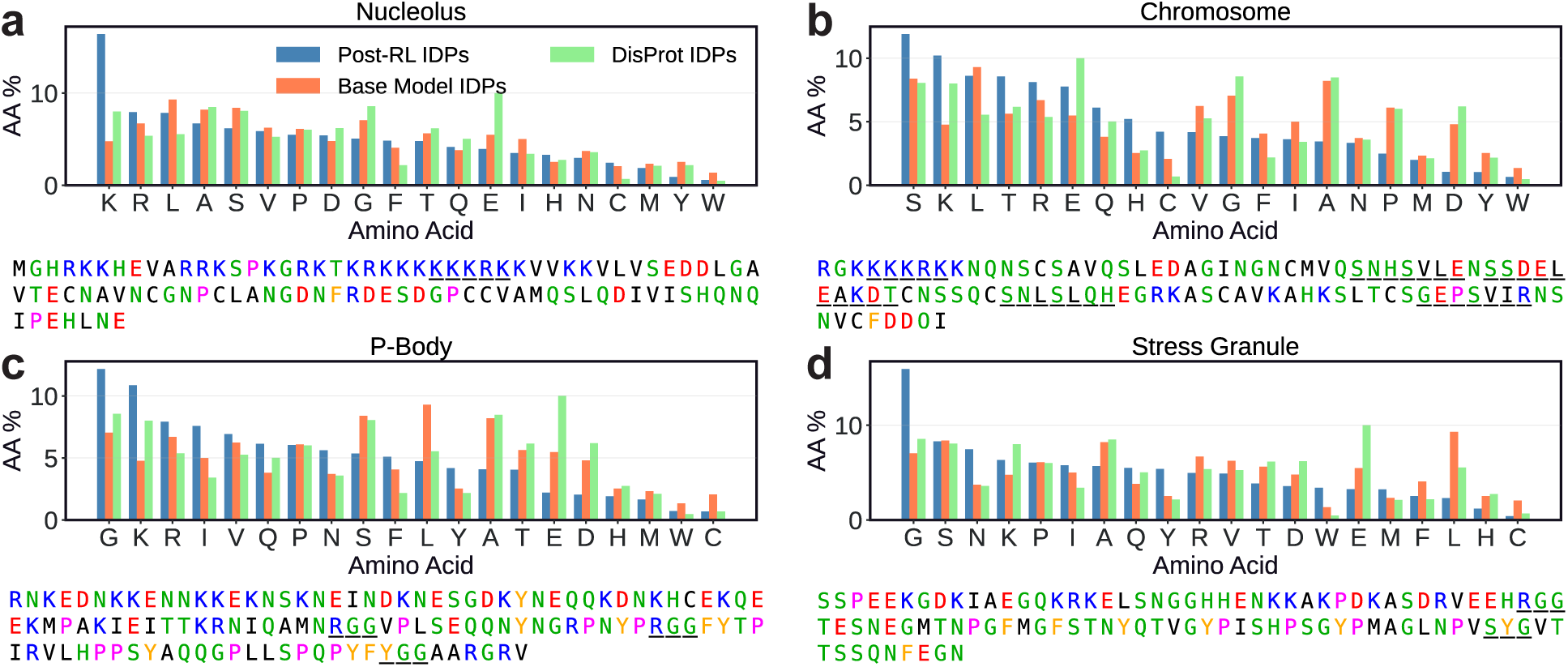
Post-trained IDiom models generate sequences which reproduce the amino acid compositional biases known to drive subcellular localization. Amino acid composition of sequences generated after post-training IDiom to optimize the ProtGPS localization score for the **(a)** nucleolus, **(b)** chromosome, **(c)** P-bodies, and **(d)** stress granules. Underneath each plot is one representative generated sequence for each compartment, with sequence features such as nuclear localization signals, post-translational modification sites, and RNA-binding motifs underlined.

Nucleolar-targeting sequences are enriched in lysine and arginine, consistent with the prevalence of positively charged nuclear localization signals and the charge-rich low-complexity regions found in nucleolar proteins [59, 60]. Sequences targeting the chromosomes are enriched in serine and threonine, consistent with the high density of phosphorylation sites characteristic of chromatin-associating proteins, which are heavily regulated by post-translational modification [61, 62]. The generated P-body targeting sequences are glycine-rich and basic (lysine- and arginine-rich), consistent with the RNA-binding motifs and arginine-glycine rich regions present in proteins that associate with RNA granules [63, 64]. Finally, stress granule-targeting sequences are similarly glycine-rich, consistent with the low-complexity, RNA-interacting motifs expected in stress granule associating proteins [65]. Representative sequences generated from each checkpoint are also shown in Figure 5, with specific sequence features such as nuclear localization signals, post-translational modification sites, and RNA-binding motifs underlined, illustrating that the global compositional shifts accompanied by the generation of specific local sequence features.

To further characterize the sequence features learned during post-training, we analyze the compartment-specific generations for sequence patterning and sequence-specific motifs. Charge segregation, quantified by *κ*, varies across compartments (Figure 6a), with nucleolus-targeting sequences showing elevated *κ* relative to all other sequences; this observation is consistent with the prevalence of charge-block architectures in nucleolar-associating IDRs [66]. In contrast, sequences targeting stress granules and P-bodies show reduced *κ* relative to baselines, consistent with the weakly charged nature of stress granule proteins [67] and the aromaticity-driven nature of P-body condensate formation [4]. In addition, we find that sequences targeting the nucleolus and chromosomes are enriched in nuclear localization signals (NLSs), as is expected for these nuclear compartments [68]. We scan generated sequences for a curated set of NLS patterns from the Eukaryotic Linear Motif (ELM) Resource [69] (patterns are provided in the Supplementary Information), and we find that a substantially higher fraction of nucleolus- and chromosome-targeting sequences contain at least one NLS compared to sequences targeting cytoplasmic compartments (Figure 6b).

**Fig. 6:**
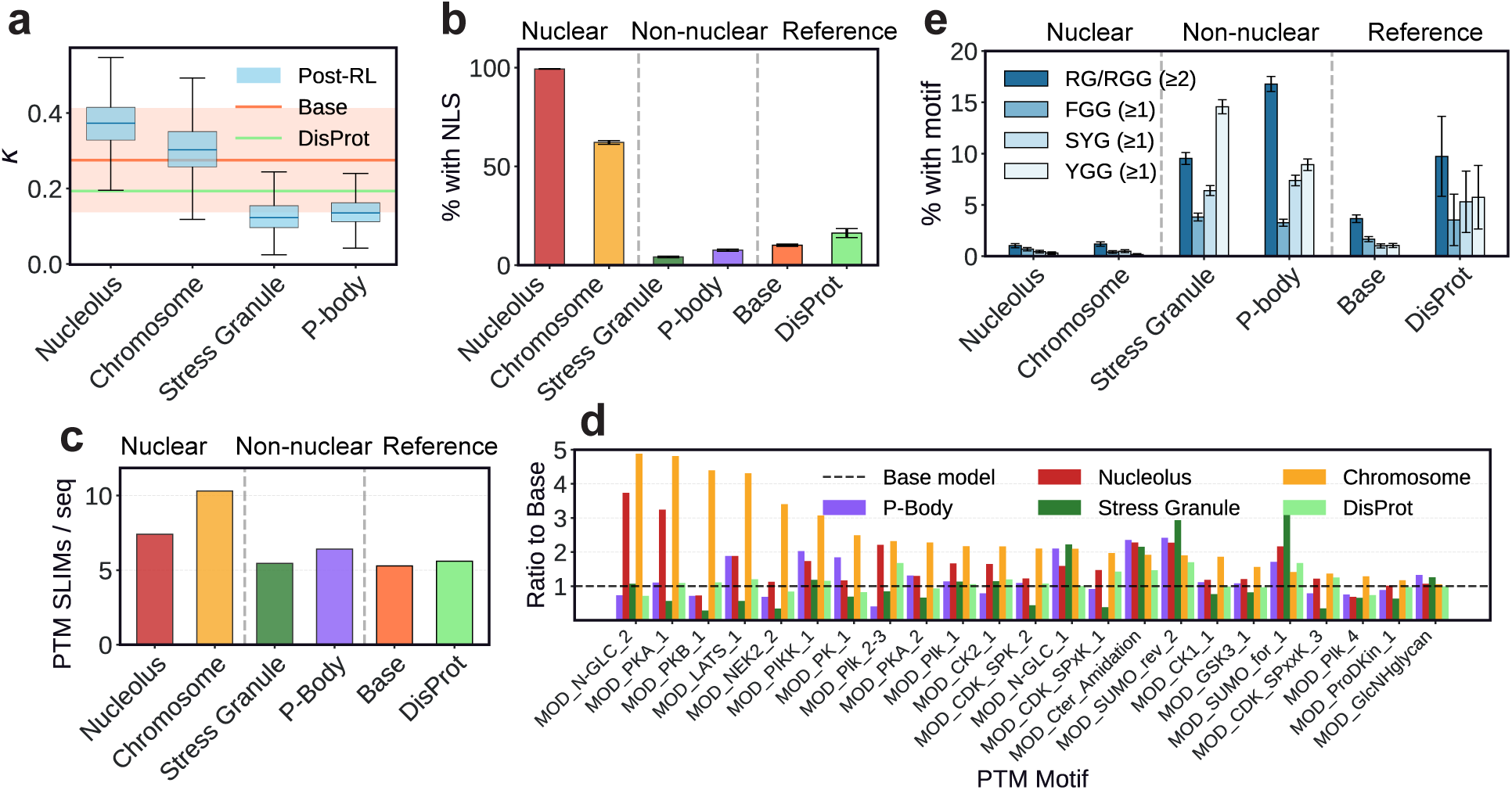
Proteins generated from post-trained IDiom models recapitulate specific sequences features which are characteristic of each target cellular compartment. **(a)** Box plot of the distribution of *κ* charge patterning values for sequences generated from IDiom checkpoints optimized for localization to the four labeled ProtGPS compartments. The horizontal orange line and band indicates the mean and standard deviation of *κ* for sequences generated from the base pre-trained model. The green line indicates the mean *κ* value of natural DisProt IDRs. **(b)** Plot of the percentage of generated sequences which contain at least one nuclear localization signal from the Eukaryotic Linear Motif (ELM) Resource. **(c)** Plot of the percentage of generated sequences which contain the RNA-interaction sequence motifs indicated in the legend. (≥ 1) indicates the presence of at least one motif within a given sequence. (≥ 2) indicates the presence of at least two motifs within 30 residues of one another within a given sequence. **(d)** Plot of the average number of unique post-translational modification (PTM) motifs from the ELM Resource per sequence for sequences generated from various model checkpoints or the DisProt IDRs. **(e)** Plot of the ratio of the number of counts of all PTM motifs from the ELM Resource (MOD) appearing in a generated sequence or DisProt IDRs, normalized to the counts within sequences generated by the base model.

Chromosome-associating proteins are heavily regulated through post-translational modifications (PTMs), and we probe whether chromosome-targeting sequences are correspondingly enriched in

PTM motifs [70]. We scan generated sequences for all 40 PTM motif classes from the ELM Resource (ELM Identifiers beginning with the MOD prefix). Chromosome-targeting sequences show a pronounced increase in MOD-motif density, with over 10 putative unique PTM sites per sequence on average, compared to approximately 5-7 sites per sequence for other compartments (Figure 6c). In Figure 6d we quantify enrichment as the ratio of motif counts in post-trained generations relative to the base model, finding that the enriched motifs span multiple kinase families, including AGC-class sites (MOD_PKA_1, MOD_PKB_1, MOD_PK_1), PIK/PIKK-associated phosphorylation sites (MOD_PIK_1, MOD_PIK_2-3, MOD_PIKK_1), acidophilic CK2 motifs (MOD_CK2_1), and proline-directed CDK-class motifs (MOD_CDK_SPK_2). This broad enrichment across kinase families is consistent with PTM-driven regulatory control being a crucial role of chromatin-interacting proteins [61, 62], and it demonstrates that post-training with a localization reward is sufficient to induce the generation of these specific sequence features.

P-bodies and stress granules are RNA-rich condensates with central roles in mRNA regulation and decay [71], and we find that post-trained sequences targeting these compartments are enriched in short RNA-interaction motifs including RG/RGG tracts, F/YGG motifs, and SYG motifs (Figure 6e). RG/RGG-rich regions are widely implicated in RNA binding and recruitment to ribonucleoprotein assemblies [63, 64], while F/YGG [72, 73] and SYG [74, 75] motifs provide aromatic sticker elements that mediate *π*-*π* and cation-*π* interactions which drive phase separation in low-complexity IDRs. The emergence of these motifs through post-training indicates that optimizing for RNA granule localization is also sufficient to induce the generation of RNA-interaction sequence grammars.

Altogether, these results demonstrate that RL post-training using the ProtGPS predictor as a reward model is able to teach IDiom the global amino acid compositions, sequence patterning, and specific motifs which are necessary for compartment-specific localization. Each of the changes we identify is biologically interpretable and consistent with the known sequence determinants of localization to or interaction with the corresponding compartment, and these features emerge without any explicit supervision. This indicates that the post-training of IDiom to optimize the ProtGPS score alone is sufficient to learn the necessary compartment-specific sequence grammars for subcellular localization of IDRs.

## Discussion

IDiom demonstrates that a protein language model trained exclusively on intrinsically disordered region sequences can faithfully capture the highly contextual evolutionary statistics of natural IDRs. The pre-trained model generates diverse sequences that recapitulate the compositional biases, charge patterning, hydrophobic decoration, and low-complexity features of natural disordered regions. Crucially, the model is able to generate highly plausible disordered sequences while remaining substantially dissimilar to the training examples. Furthermore, IDiom learns in-context and is able to generate IDRs conditioned on flanking sequence context that are more similar to the natural IDRs that exist within that flanking context, as demonstrated by the DisProt context-prompted IDRs and the NPM1 case study.

Beyond generation from the base pre-trained model, we demonstrate that IDiom can be post-trained using reinforcement learning to optimize arbitrary external reward functions. While here we demonstrate this using ProtGPS as the reward signal for subcellular localization, the post-training approach is general. IDiom can be combined with any objective, such as sequence-based predictors of conformational ensembles [76], intermolecular interactions [45], phase behavior [77], as well as machine-learned predictors of experimentally characterized biological function such as transcriptional activity [78, 79] or direct binding affinity [80]. Post-training can also be applied directly to experimental data, using approaches such as direct preference optimization [54, 81] or energy rank alignment [82, 83]. Such approaches would enable iterative optimization of the generative model as new experimental measurements become available. Due to the often modular nature of IDRs, a designed IDR could be inserted or appended to a full length protein to tune localization, phase behavior, or signaling properties without redesigning the folded domains. Additionally, the prevalence of short IDRs in our training data (Figure 1e) also makes IDiom well-suited for the design of disordered peptides for therapeutic applications. Finally, IDiom offers interesting opportunities for shrinking proteins containing functional IDRs, an important objective in protein delivery [84].

IDiom can also enable the automated discovery of evolutionarily and biologically important sequence features within IDRs. In the examples with the ProtGPS reward model, we manually identified sequence features to analyze in the post-trained sequences. Future work applying techniques such as feature learning with sparse autoencoders [85] would enable automatic identification of prominent sequence features learned by IDiom. While prior work has studied the features learned by protein language models trained on full length sequences, intrinsically disordered regions are poorly resolved in those learned feature spaces [86]. In contrast, sparse autoencoders trained on IDiom representations would be free from folded domain features, allowing for more precise identification of sequence grammars that underlie intrinsically disordered regions. When applied to post-trained models, this approach could further reveal the function-specific features that underlie biological function, such as subcellular localization.

Intrinsically disordered regions play central roles in cellular processes such as gene regulation, subcellular compartmentalization, and signaling, yet they have remained largely inaccessible to rational design. IDiom provides a generative framework for disordered sequence design that can be steered towards specific functional objectives through post-training. Combined with high-throughput experimental assays and machine learned reward models trained on experimental data, this platform offers a path toward the systematic design and engineering of intrinsically disordered proteins and regions, opening new avenues in synthetic biology, targeted therapeutics, and the design of programmable condensates.

## Methods

### Data Curation

We curate our dataset of intrinsically disordered regions from the AlphaFold Database (AFDB), version 4. First, we use MMseqs2 to cluster the 214M AFDB sequences at 90% identity and 80% coverage (other MMseqs2 parameters below). Next, we follow the method of [32] to determine the sequences and locations of pLDDT-based IDRs within the MMseqs2 cluster representative proteins. In this method, we first apply a 15 residue-wide averaging filter to the pLDDT values. Next, we mark residues with pLDDT > 80 as folded, pLDDT < 70 as disordered, and 70 < pLDDT < 80 as gap regions. Folded and disordered regions with a length shorter than ten residues are reclassified as gaps. If a gap region is flanked by two disordered regions, or if it is N- or C-terminal and is adjacent to a disordered region, we relabel it as disordered. All other gap regions are relabeled as folded. Each record within the dataset corresponds to a different IDR, and any given protein may yield ≥ 1 IDR. IDRs which are located in proteins whose full length is greater than 512 residues, IDRs shorter than 30 residues long, and sequences whose entire length is low-pLDDT are discarded. This curation process yields 37M IDRs and their associated N- and C-terminal flanking contexts.

### Sequence Clustering and Percent Identity Characterization

We use MMseqs2 to cluster sequences in the AlphaFold Database at 90% identity and 80% coverage. MMseqs2 was run using the following command: mmseqs linclust --min-seq-id 0.9 --cov-mode 0 -c 0.8 --cluster-mode 2.

We also use MMseqs2 to identify the sequence identity of generated sequences with respect to the training set. Specifically, we compare generated IDR and IDP sequences against the 37M AFDB training set IDRs without their surrounding context. MMseqs2 was run with the following command to search the generated sequences against the AFDB training set IDRs: mmseqs search --max-seqs 1 -e 1e3 --min-seq-id 0.0 -c 0.0. For each generated sequence, this command returns the sequence identity of the closest match in the training set, which is plotted in Figure 1.

### Tokenization

We tokenize protein sequences using a simple alphabet in which each amino acid is represented by a single token. We introduce the tokens <N>, <C>, and <I> to denote the N-terminal flanking context of an IDR, the C-terminal flanking context, and the IDR span itself, respectively. We additionally add the standard beginning-of-sequence <BOS> and end-of-sequence <EOS> tokens to all sequences, and we pad all sequences to the maximum length of 512 tokens using the <PAD> token. The total size of the alphabet we use is 27 tokens (20 amino acids, <N>, <C>, and <I>, and <BOS>, <EOS>, <PAD>, and <MASK>.

### Data Augmentations

To process the curated AFDB IDRs for model pre-training, we transform the protein sequences into a fill-in-the-middle format. We prepend the token <N> to any N-terminal context of the IDR, prepend to the IDR itself, and prepend <C> to any C-terminal context of the IDR. We then rearrange the sequence in the order <N><N-terminal context><C><C-terminal context><IDR span>. An example sequence is <N>MEDS..HLVA<C>SVED..RKSLVEED..KGPS.

We augment the data with intrinsically disordered proteins by duplicating the set of IDRs and removing their N- and C-terminal flanking contexts. An example sequence from this data augmentation is <N><C>VEED..KGPS. In all, this produces 74M sequences for training (37M IDRs and 37M IDPs).

### Model Architecture

IDiom is a 12-layer decoder-only Transformer with 14 attention heads and a hidden dimension of *d*_model_ = 896. The model employs pre-LayerNorm and utilizes the SwiGLU non-linearity [87] in all feedforward networks. The feedforward network expansion ratio is 8/3, resulting in a total of 122M trainable parameters. Positional information is encoded using Rotary Position Embeddings (RoPE) [88]. The model processes sequences with a maximum length of 512 tokens, and shorter sequences are padded to this length. Multi-head attention is computed using Flash Attention [89].

### Pre-training

We pre-train IDiom on the aforementioned 74M sequences. The data are randomly split into 99% training, 0.5% validation, and 0.5% test sets. All sequences are padded or truncated to a fixed length of 512 tokens. The model is trained autoregressively using next-token prediction.

Training is performed using the AdamW optimizer with a learning rate of 4.0×10*^−^*^4^ and no weight decay. The learning rate schedule uses a linear warmup over the first 3,000 steps, followed by cosine annealing decay to a minimum of 4.0 × 10*^−^*^5^ (10% the initial learning rate) over 250,000 total training steps. The global batch size is 1,024 (Distributed Data Parallel training with 128 sequences per GPU across 8 NVIDIA H100 GPUs), with no gradient accumulation. We use a cross-entropy loss while ignoring contributions from pad or mask tokens. Training is performed in mixed precision (fp32/bfloat16) and no gradient clipping is applied. Model validation is performed every 25,000 training steps on the validation set, and training is run for 250,000 optimizer steps. The training and validation curves are presented in the Supplementary Information.

### Sequence Generation

We generate IDR sequences from IDiom using autoregressive decoding. At each position *t* in the sequence, we compute the model logits *z_t_*and convert them to a probability distribution via *p*(*x_t_*|*x_<t_*) = softmax(*z_t_/T*), then sample the next token from this full categorical distribution over the vocabulary. We use a fixed sampling temperature of *T* = 1.0 for all generations

Sequence generation supports both prompted and unprompted modes. For prompted generation, the N- and C-terminal flanking contexts are provided as the prompt in fill-in-the-middle format: <N><N-terminal context><C><C-terminal context>, and the model generates the IDR span autoregressively following the token. For unprompted generation of fully disordered proteins, the prompt consists of only the three special tokens without any flanking context: <N><C>, and the model generates the disordered sequence. In both cases, generation terminates upon sampling the <EOS> token or upon reaching the maximum sequence length of 512 tokens.

### Post-training

We fine-tuned IDiom using Group Relative Policy Optimization (GRPO) with the Decoupled Advantage Policy Optimization (DAPO) modification. During GRPO training, for each batch of sequence prompts, the model generates multiple completions per prompt (group size = 8). Rewards are computed for each generated sequence using a reward function that combines three rewards: 1. A quadratic reward around a target ProtGPS score of 0.9 for the desired target compartment, 2. A quadratic reward around a target sequence length of *L*_target_ = 100 residues, and 3. A quadratic reward around a target sequence Shannon entropy of *H*_target_ = 2.7 nats to prevent diversity collapse.

Within each group, advantages are computed as normalized relative rewards: 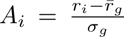, where 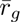 and *σ_g_* are the group-wise mean and standard deviation. The GRPO loss combines a clipped policy gradient term (PPO-style clipping with *ɛ*_clip_ = 0.2) weighted by advantages, and a KL-divergence penalty L_KL_ = *β*_KL_*D*_KL_(*p*_ref_||*p*_current_) with strength *β*_KL_ = 0.02. The total loss is optimized per-token on generated completions only (any prompt tokens are masked).

Post-training is performed with a learning rate of 5×10*^−^*^6^, the AdamW optimizer, and a global batch size of 8, for 1,500 optimizer steps. Post-training was performed separately for all 12 target cellular compartments in ProtGPS, although we only analyze the sequences generated from checkpoints trained for localization to the nucleolus, chromosomes, P-bodies, and stress granules.

### Structure Prediction

We use Colabfold [37] to perform AlphaFold2 structure predictions on generated sequences and determine the predicted local distance difference test (pLDDT) values.

### Sequence Analysis

We use the Sparrow package [90] to calculate sequence-level metrics such as charge patterning *κ*, sequence hydropathy decoration, sequence complexity (SEG), and fraction of charged residues. Short linear motif (SLIM) analysis is performed with a regular expression search using the SLIM regular expressions from the Eukaryotic Linear Motif Resource [69]. The specific regular expressions are presented in the Supplementary Information

### Visualization

Protein structure visualization is performed with PyMOL [91].

## Data and Code Availability

The code used for model pre-training, sequence generation, and post-training is available on Github: Code (https://github.com/rotskoff-group/idiom).

The pre-training datasets and pre- and post-trained model checkpoints are available on Hugging Face: Datasets (https://huggingface.co/datasets/jxliu2/idiom-datasets) Models (https://huggingface.co/jxliu2/idiom).

Information regarding these artifacts is provided in the Supplementary Information.

## Acknowledgements

J.X.L. acknowledges support from the National Institute of Health T32 award number T32HL094274. S.I. acknowledges support from the National Science Foundation Graduate Research Fellowship Program and the Shoucheng Zhang Graduate Fellowship. F.H. acknowledges support from the Stanford Center for Molecular Analysis and Design Fellowship. Research reported in this publication was supported by the National Institute of General Medical Sciences of the National Institutes of Health under award number 1R35GM159834-01 (G.M.R.) and 5R35GM130332 (A.R.D.). The content is solely the responsibility of the authors and does not necessarily represent the official views of the National Institutes of Health. The authors acknowledge the use of the Stanford Sherlock compute cluster, and the authors acknowledge insightful discussions with Prof. Mikko Haataja.

## Supplementary Information

### Datasets

Here we describe the datasets provided on HuggingFace:

Datasets (https://huggingface.co/datasets/jxliu2/idiom-datasets)

Below, we describe the data files under idr_datasets/training_sequences:

- AFDB_IDR_90_reps.fasta contains the 53M cluster representatives after the initial 214M full length AFDB protein sequences are clustered at 90% identity, 80% coverage.
- AFDB_IDR_90_alldata.h5 contains 73M IDRs as extracted from the AFDB according to the Tesei logic [32] (see Methods), and after filtering for IDRs belonging to the 53M cluster representatives identified in AFDB_IDR_90_reps.fasta. This HDF5 file contains the following keys: <KeysViewHDF5 [’accession_ids’, ‘full_avg_plddt’, ‘full_length’, ‘full_-seq’, ‘idr_end’, ‘idr_length’, ‘idr_plddt’, ‘idr_start’, ‘idrs’]>.
- AFDB_IDR_90_FIM_512.h5 is created from AFDB_IDR_90_alldata.h5 by filtering out IDRs whose full length sequences are longer than 512 residues. We also find that ∼ 1/3 of records in AFDB_IDR_90_alldata.h5 are fully low-pLDDT sequences, and we filter out those sequences because we find that they are not representative of intrinsically disordered proteins. We only keep sequences with both low- and high-pLDDT regions. We hypothesize that sequences which are fully low-pLDDT are due to AlphaFold2’s poor confidence in sequences which are not similar to those seen during training, rather than because they are fully intrinsically disordered proteins. For the remaining 37M IDRs, we apply the fill-in-the-middle (FIM) transformation as well as IDP data augmentation as mentioned in the Methods, and place those records into AFDB_IDR_90_FIM_512.h5. We note that we represent the <N>, <C>, and tokens with 1, 2, and 3, respectively, in this HDF5 file as well as in the codebase. This is the final file used for the precompute and pre-training steps.
- AFDB_IDR_90_FIM_512_full.fasta contains the 37M full length sequences (in correct order, not FIM-transformed) contained in AFDB_IDR_90_FIM_512.h5. The fasta header contains_IDR_X-Y where X and Y are the 1-indexed indices of the start and end (inclusive) of the intrinsically disordered region.
- AFDB_IDR_90_FIM_512_idrs.fasta contains only the sequences of the 37M intrinsically disordered regions in AFDB_IDR_90_FIM_512_full.fasta, without their surrounding context.

We also provide several datasets of sequences generated by our model under idr_datasets/generated_-sequences. All generated sequences are provided in FASTA format along with their corresponding autoregressive model log (pickle format).

- Generated IDPs: 100,000 unprompted intrinsically disordered proteins.
- Generated IDRs: 101,700 intrinsically disordered regions generated using 1,017 DisProt flanking contexts prompts (100 generated IDRs per prompt).
- Generated NPM1 IDRs: 100,000 sequences generated using the NPM1 flanking context as the prompt (UniProt: P06748).
- Generated ProtGPS Sequences: 10,000 IDPs generated from post-trained checkpoints. Post-training was done to optimize ProtGPS localization scores for the four target compartments: chromosome, nucleolus, P-body, and stress granule.

### Models

Here we describe the model checkpoints and other files provided on HuggingFace:

Models (https://huggingface.co/jxliu2/idiom)

Below, we describe the directories under idiom/:

- base/ contains the checkpoint of our pre-trained base IDiom model, along with its configuration files.
- post_trained/protgps_reward/ contains the checkpoints of IDiom post-trained via reinforcement learning using the ProtGPS reward model, one checkpoint per target compartment. In this paper, we analyzed results for 4 compartments: the nucleolus, stress granules, P-bodies, and chromosomes. However, post-training runs were conducted for all 12 ProtGPS compartments (chromosome, nucleolus, nuclear speckle, nuclear pore complex, P-body, PML body, post-synaptic density, stress granule, Cajal body, RNA granule, cell junction, and transcriptional condensate). We leave analysis of the remaining compartments to future work.
- protgps/ contains the ProtGPS reward model used during reinforcement learning post-training.
- data/ contains auxiliary files used during training and inference.

### Secondary Structure Metric Analysis

Here we present analysis of secondary structure metrics for training, generated, DisProt, and CATH sequences. Secondary structure was assigned per residue using the dictionary of secondary structure of proteins (DSSP) algorithm as implemented in MDTraj [92]. Secondary structure content was then defined as the sum of the mean *α*-helical and mean *β*-sheet fractions across all residues. Fig. S1 shows histograms of the average secondary structure content of 100 randomly chosen sequences from the various training, generated, DisProt, and CATH sets of proteins.

**Fig. S1:**
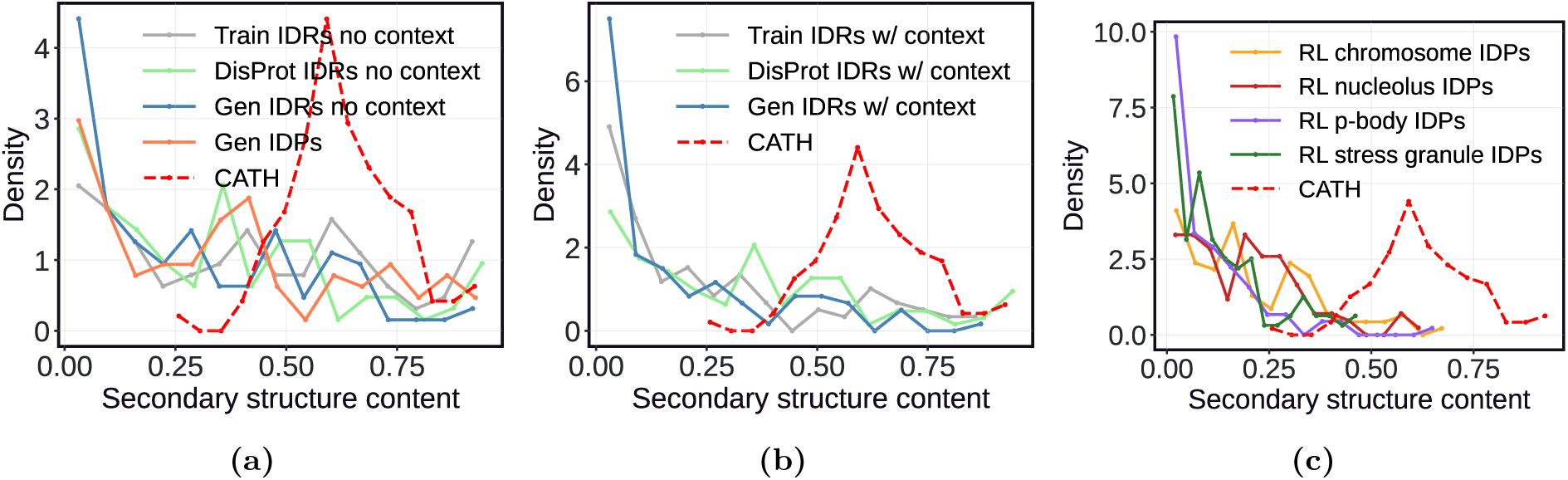
Secondary structure content analysis. Histograms of of the average secondary structure content (*α* + *β*) for the AF2-predicted structures of 100 randomly chosen sequences from the following sets of sequences: **(a)** Secondary structure content of training IDRs, generated IDRs, DisProt IDRs, and CATH sequences, with their structures predicted with surrounding context included. **(b)** Secondary structure content of training IDPs, generated IDPs, DisProt IDPs, and CATH sequences, with their structures predicted without their surrounding context. **(c)** Secondary structure content of IDPs generated from post-trained IDiom checkpoints and CATH sequences, with their structures predicted with surrounding context included.

### Disorder Predictions

Here, we present orthogonal disorder predictions from Metapredict v3 [93] and IUPred3 [94]. For both predictors, a higher value represents a higher propensity towards disorder.

**Fig. S2:**
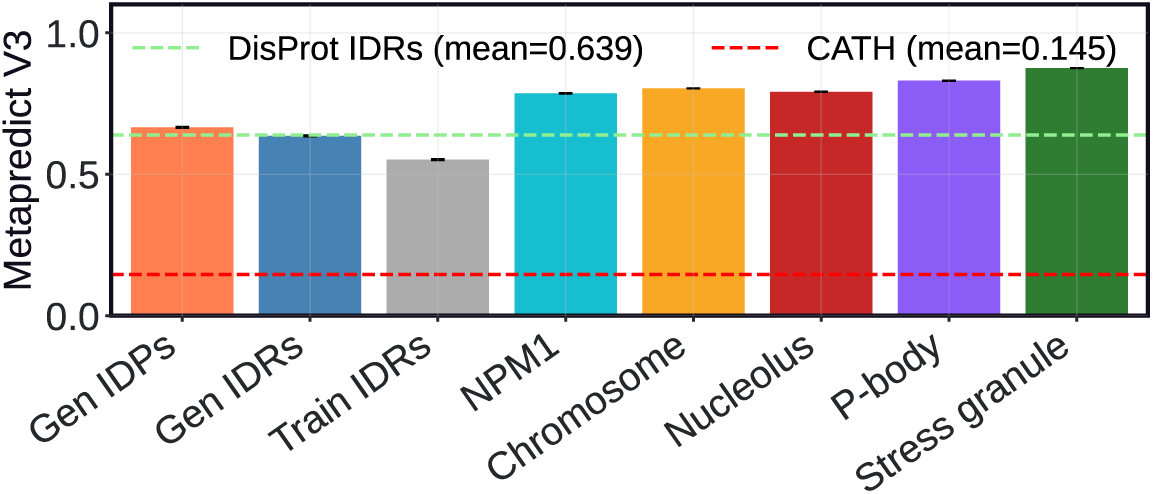
Disorder predictions from Metapredict V3. Higher values correspond to higher propensity towards disorder. The horizontal green and red dashed lines correspond to the predicted Metapredict V3 values for 1,017 DisProt IDRs and 1,000 CATH sequences, respectively. The bars correspond to predicted Metapredict values for 10,000 sequences generated from IDiom for each condition, as well as 10,000 training sequences. The sequences generated from IDiom include unprompted IDPs, DisProt-prompted IDRs, NPM1 IDRs, and IDPs generated after post-training for localization to the chromosomes, nucleolus, P-bodies, and stress granules.

**Fig. S3:**
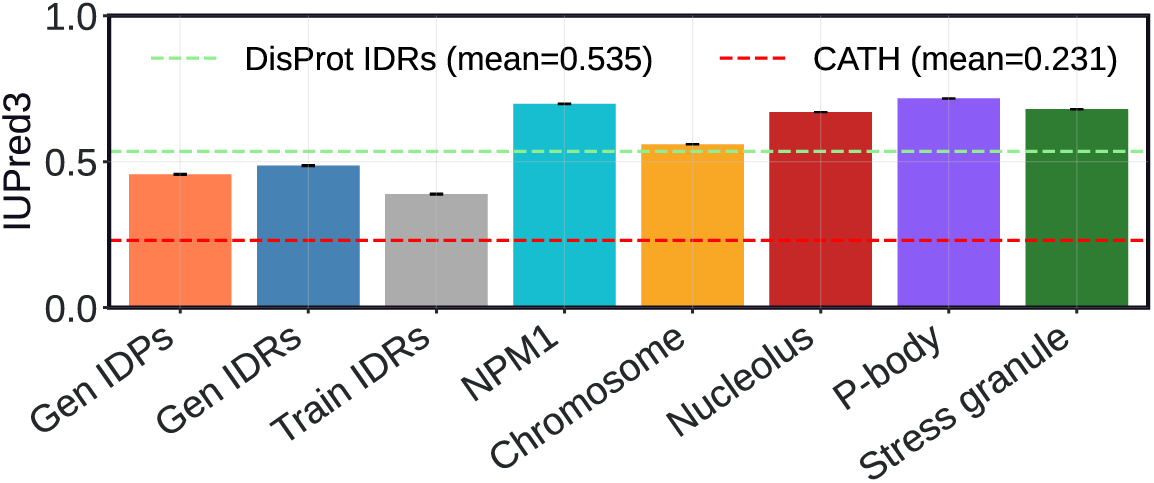
Disorder predictions from IUPred3. Higher values correspond to higher propensity towards disorder. The horizontal green and red dashed lines correspond to the predicted IUPred3 values for 1,017 DisProt IDRs and 1,000 CATH sequences, respectively. The bars correspond to predicted IUPred3 values for 10,000 sequences generated from IDiom for each condition, as well as 10,000 training sequences. The sequences generated from IDiom include unprompted IDPs, DisProt-prompted IDRs, NPM1 IDRs, and IDPs generated after post-training for localization to the chromosomes, nucleolus, P-bodies, and stress granules.

### ESM3 Comparison

Here we present comparison plots between sequences generated by IDiom and ESM3, using the same 1,017 DisProt flanking domain prompts. ESM3 sequences are generated using iterative decoding. A total of 1,000 sequences are sampled for each prompt. As ESM3 consists of a bidirectional transformer architecture, the length of the generated IDRs is fixed at the length of the ground truth IDR. The number of decoding steps, i.e. forward passes until the sequence is fully unmasked, is set to be the minimum of 20 and the ground truth IDR length for each prompt. Tokens are sampled with a temperature of 1.0, and all other default inference hyperparameters for ESM3 are used. We find that compared to IDiom, ESM3-generated IDRs are extremely low-complexity sequences, with a peak in the SEG complexity distribution around 0.5 (Figure S4). We show three example ESM3-generated IDRs below, in red:

**Fig. S4:**
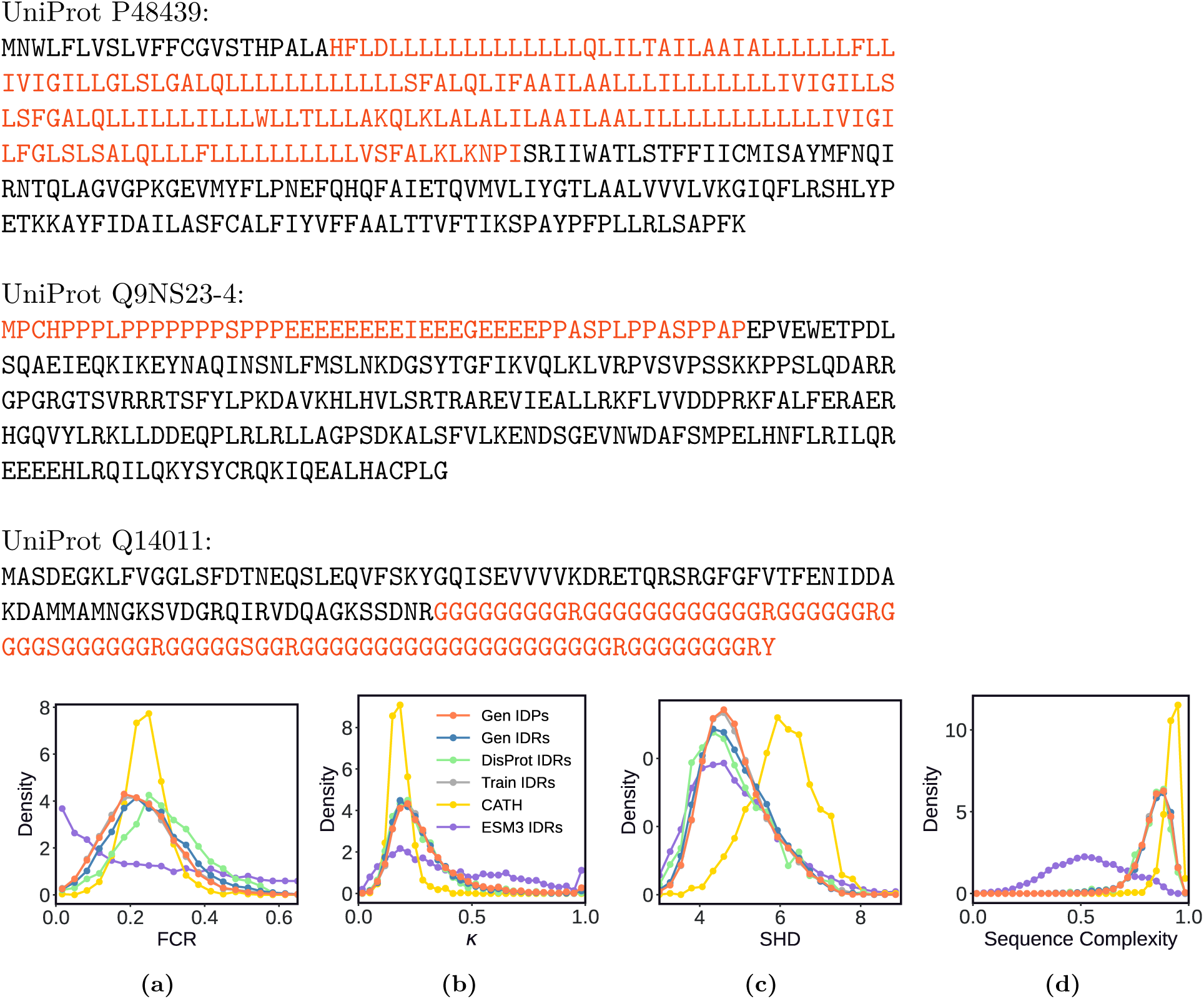
Comparison between ESM3-generated IDRs and IDiom-generated IDRs. **(a)–(d)** Distributions of various sequence metrics for sequences generated from IDiom versus ESM3. Training set IDRs, natural DisProt IDRs, and folded CATH domains are shown as well. **(a)** Fraction of charged residues (FCR). **(b)** Charge patterning *κ* parameter. **(c)** Sequence hydropathy decoration (SHD). **(d)** Sequence complexity quantified by the SEG algorithm.

### Short Linear Motifs from the Eukaryotic Linear Motif Resource

Here, we list the short linear motifs from the ELM Resource which we scan for, for nuclear localization signals (NLSs) as well as for post-translational modification (PTM) sites (ELM Identifier: MOD).

**Table 1:**
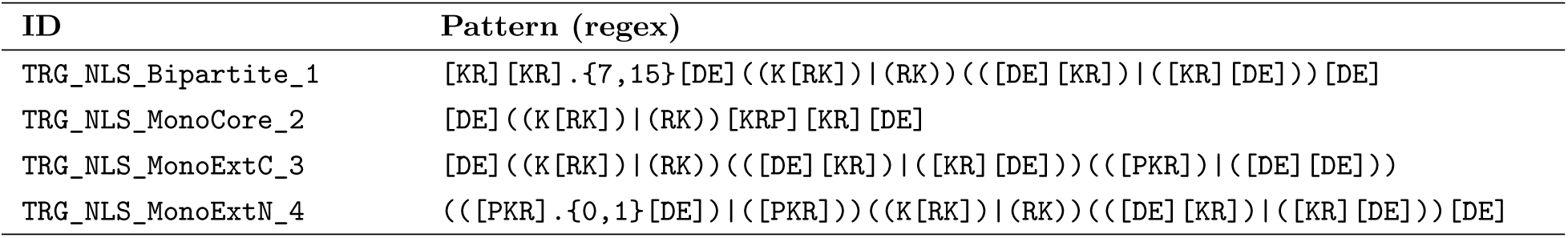
ELM NLS motifs and their corresponding regex patterns.

#### Nuclear Localization Signals

The regular expressions of the 4 NLSs we consider are:

#### Post Translational Modification Motifs

The regular expressions of the 40 PTM MOD sites we consider are:

**Table 2:**
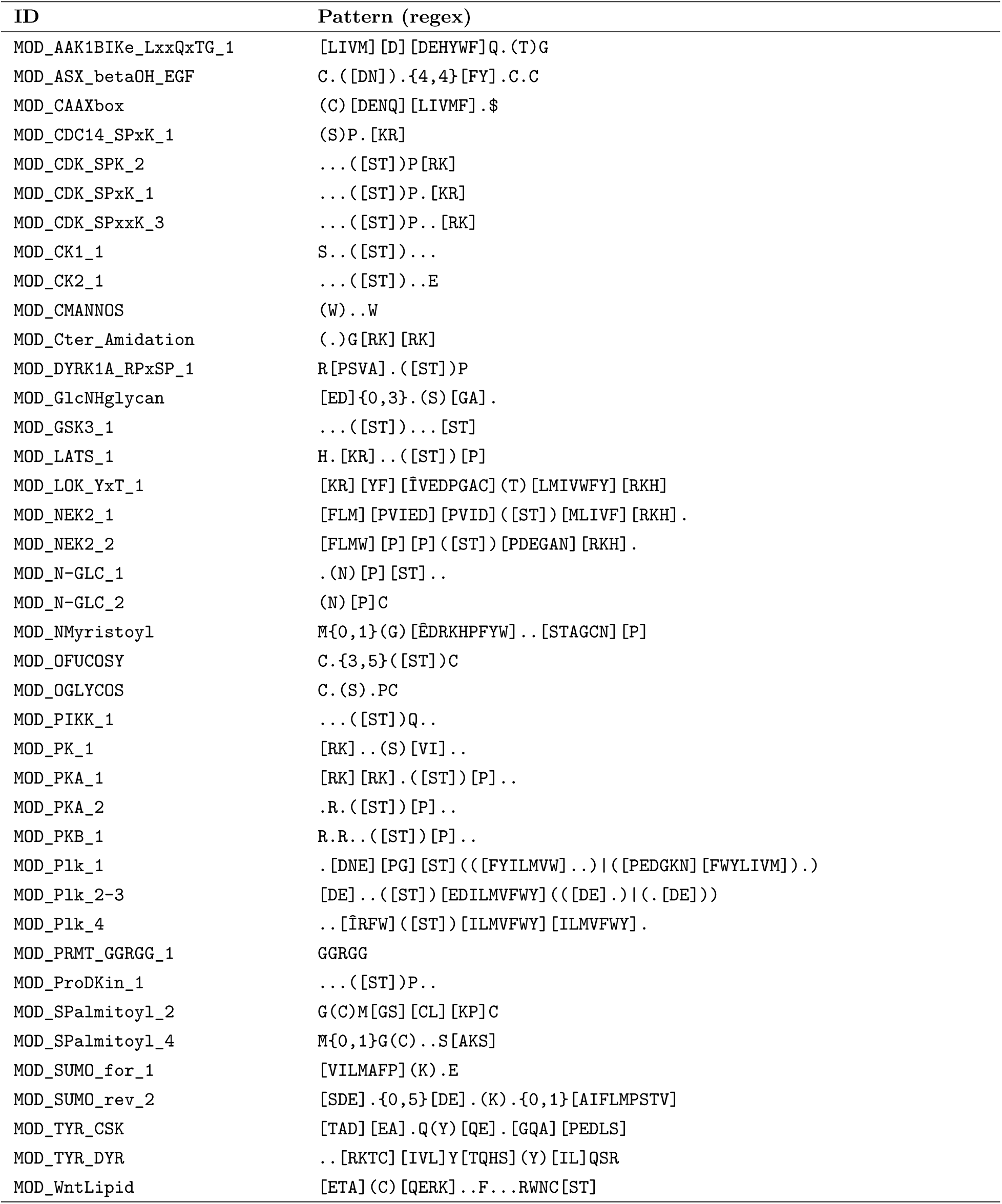
ELM MOD motifs and their corresponding regex patterns.

### Training Curves

Here, we present additional training curves from pre-training as well as post-training.

**Fig. S5:**
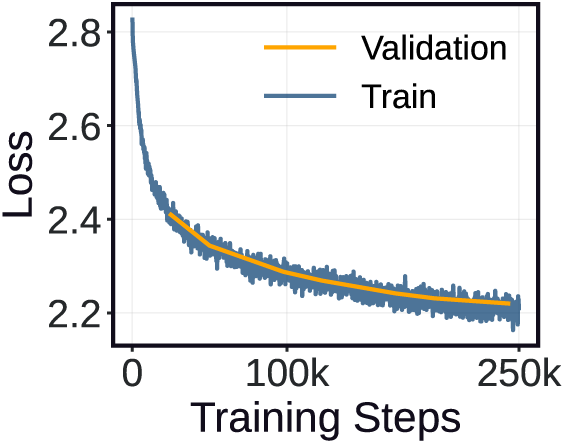
Pretraining loss curves. Training and validation losses vs optimizer steps during pre-training. The final training loss is 2.19. The final validation loss is 2.22.

**Fig. S6:**
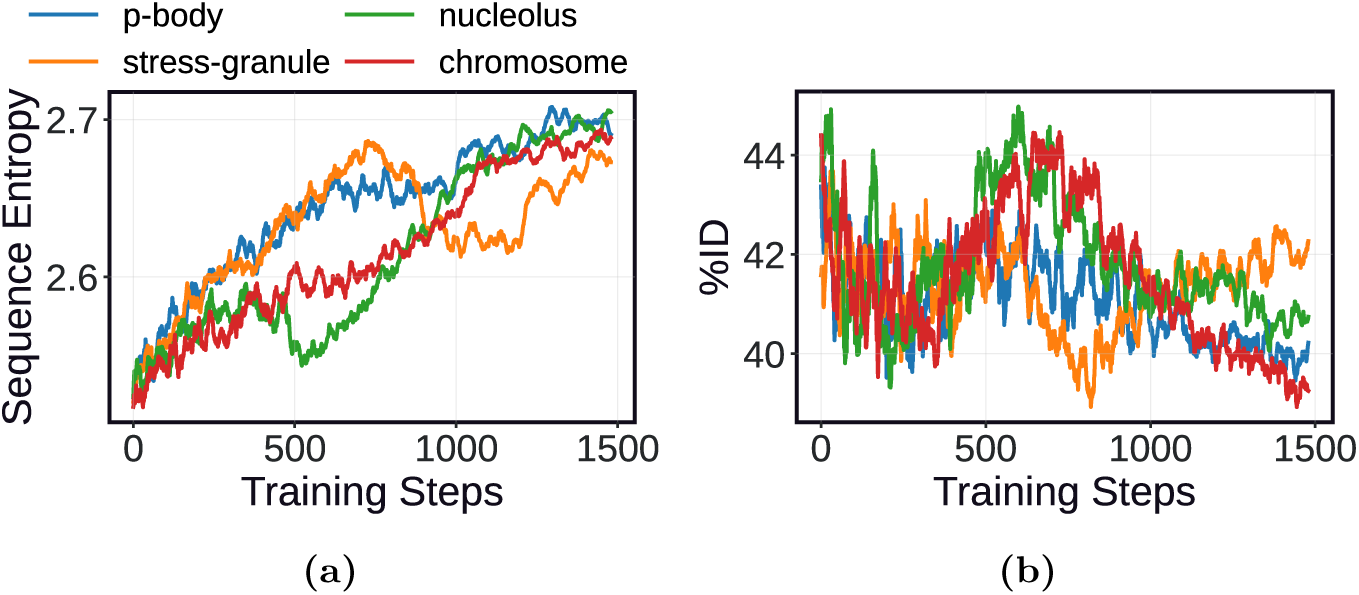
Additional post-training curves with the ProtGPS reward model. **(a)** Shannon entropy vs. training steps (target *H* = 2.7). **(b)** %ID within a generated batch vs training steps (no target value).

